# RNA-Seq analysis of genes affected by Cyclophilin A/DIAGEOTROPICA (DGT) in tomato root development

**DOI:** 10.1101/2020.08.29.272567

**Authors:** Maria G. Ivanchenko, Olivia R. Ozguc, Stephanie R. Bollmann, Valerie N. Fraser, Molly Megraw

**Author notes:** Co-first authors (Maria G. Ivanchenko, Olivia R. Ozguc).

## Abstract

Cyclophilin A/DIAGEOTROPICA (DGT) has been linked to auxin-regulated development in tomato and appears to affect multiple developmental pathways. Loss of DGT function results in a pleiotropic phenotype that is strongest in the roots, including shortened roots with no lateral branching. Here, we present an RNA-Seq dataset comparing the gene expression profiles of wildtype (‘Ailsa Craig’) and *dgt* tissues from three spatially separated developmental stages of the tomato root tip, with three replicates for each tissue and genotype. We also identify differentially expressed genes, provide an initial comparison of genes affected in each genotype and tissue, and provide the pipeline used to analyze the data. Further analysis of this dataset can be used to gain insight into the effects of DGT on various root developmental pathways in tomato.

## Introduction

The tomato (*Solanum lycopersicum*) cyclophilin DIAGEOTROPICA (DGT) has been linked to auxin-regulated development through identification of the gene affected by the *diageotropica* (*dgt*) mutation^1,2^. Tomato *dgt* mutants are auxin-resistant and display a pleiotropic phenotype that includes slow gravitropic response, lack of lateral root initiation, altered vascular development, reduced ethylene production in response to auxin, reduced apical dominance, impaired shoot and root growth, reduced fertility, and impaired fruit growth^3–11^. DGT likely interacts with auxin transport and signaling in a complex manner. For instance, DGT has been shown to negatively regulate auxin efflux via PIN-FORMED (PIN) transporters by altering subcellular localization of PINs and expression of some *PIN* genes^9^. *DGT*, in turn, is downregulated by auxin at the root tip^8^, suggesting functional feedback between DGT, auxin, and PINs. Through targeted gene expression quantification with RT-PCR and northern blots, DGT has been demonstrated to affect expression of a number of other auxin-related genes in addition to *PIN* genes. The *dgt* mutation reduces auxin-induced expression of genes encoding certain 1-aminocyclopropane-1-carboxylic acid synthases (ACCSs; key ethylene biosynthesis regulatory enzymes), SMALL AUXIN UPREGULATED RNA (SAUR) genes, and several members of the auxin-regulated *Aux/IAA* gene family in a tissue- and developmental stagespecific manner^4,6,10–14^. However, full transcriptomic profiling of tomato *dgt* mutants in different developmental zones has not been performed. Given the complex role of DGT in auxin pathways, an exploration into the widespread effects of DGT on the transcriptome may provide valuable insights into its potentially extensive and multifaceted role in development.

In this study, we perform a global analysis of gene expression in *dgt* roots that compares the root meristem, elongation zone, and differentiation zone in wildtype (‘Ailsa Craig’) and *dgt* tomato plants. The root tip provides an excellent system for studying development-related plant gene expression because cell division, elongation, and maturation are not only temporally but also spatially separated in this growth region; this allows for anatomical dissection and analysis of specific developmental zones, such as the meristem, elongation zone, and differentiation zone. The root tip is also most appropriate for this study because the *dgt* phenotype is the strongest in the root tip and has been characterized morphologically in the root tip^7,8^. We have previously performed histological analyses of tomato root tips including the meristem, elongation zone, and maturation zone, and demonstrated a decrease in length and number of cells of the *dgt* meristem and elongation zone^8^, whereas the initiation of lateral root primordia was abolished in the *dgt* root maturation zone^7–9^. Here, we present an RNA-Seq dataset containing raw reads and abundance estimates for three replicates in each zone and genotype, the pipeline used for analysis, and an initial exploration of expressed and differentially expressed genes in each developmental region that can be used to guide future investigations.

## Materials and Methods

### Plant material and growth conditions

Three biological replicates for each tissue and genotype were performed. Seven-day old tomato (*Solanum lycopersicum*) wildtype (WT) and *dgt^1-1^* plants in the ‘Ailsa Craig’ background^7^ were used. Seeds were sterilized in 20% commercial bleach for 30 min and rinsed four times for 10 min with sterile water. Sterilized seeds were vernalized at 4°C for 2 days to ensure even germination and then planted on media containing 0.2× Murashige and Skoog basal medium with vitamins (PhytoTechnology; http://www.phytotechlab.com), 1% sucrose, 10 mM MES buffer pH 5.7, and 0.8% agar. Seeds were germinated in Magenta boxes (16 seeds per box) in a growth chamber at 21 °C under long day (16h light, 8h dark) conditions and light intensity as in Ivanchenko et al. (2013). Root samples were dissected under DIC optics at 4× objective as in Ivanchenko et al. (2006). On average, 50–100 root portions were collected per biological replicate. The meristem was dissected between the root tip and the root portion where tissue becomes more transparent. The elongation zone was collected from the proximal meristem border and the first hair bulge, and approximately 1 cm portions were collected above the first hair bulge for the differentiation zone.

### RNA isolation and RNA-Seq

Tissue samples were collected in Plant RNA Reagent (Life Technologies; http://www.lifetechnologies.com) on ice, and total RNA was prepared using the RNeasy kit (Qiagen; http://www.qiagen.com) according to manufacturer’s recommendations. The RNA pellets were dried in 1.7 mL centrifuge tubes and solubilized in 178 *μ*L 1X RNA Secure Reagent (Ambion; http://www.lifetechnologies.com) preheated at 65°C, and incubated at 65°C for 10 min, mixing by pipetting a few times. Then 20 *μ*L 10X DNase I buffer and 2 *μ*L RNase-free DNase I (Ambion; http://www.lifetechnologies.com) were added to each tube, and tubes incubated for 10 min at 37°C. 700 *μ*L RLT buffer was added (to which 7 *μ*L 2-Mercaptoethanol was freshly added) to each sample, which were then mixed by vortexing. 500 *μ*L ethanol was then added and samples were mixed again by vortexing. Each sample (2 × 700 *μ*l) was applied to an RNeasy Mini spin column from the RNeasy kit and RNA cleanup performed following the manufacturer’s instructions. Each sample was eluted with 30 *μ*L nuclease-free ultrapure H_2_O, and the RNA concentrations were measure using a NanoDrop 1000 spectrophotometer (Thermo Fisher; https://www.thermofisher.com/).

### Illumina Sequencing of RNA

RNA-Seq library preparation and sequencing were performed at the Oregon State University Center for Genome Research and Biocomputing. Libraries were prepared using the TruSeq RNA Sample Prep Kit v2 (Illumina; https://www.illumina.com/) and sequenced as single-end 51 bp reads on the Illumina HiSeq 2000 using a total of two lanes.

### Gene alignment and expression level analysis

Sequencing produced 5-8 FASTQ files per replicate, which were merged into a single FASTQ file per replicate. Reference transcripts were extracted and preprocessed from the NCBI *Heinz 1706* genome sequence^15^ using RSEM’s^16^ (RNA-Seq by Expectation Maximization, version 1.3.1) rsem-prepare-reference function. Using RSEM, raw sequence reads were then aligned to the reference transcript sequences and estimated transcripts per million (TPM) values were calculated using default parameters. Resulting gene-level estimates from biological replicates were merged into a single input matrix and EBSeq (version 1.26.0) was then used to test for differential expression between *dgt* and WT for each root-tip zone. Raw RNA-Seq reads, abundance estimates, and differential expression analysis are publicly available in the Sequence Read Archive (SRA; https://www.ncbi.nlm.nih.gov/sra) and Gene Expression Omnibus (GEO; https://www.ncbi.nlm.nih.gov/geo/).

### Data Visualization

As a preliminary analysis to guide future investigations, a basic comparison of presence/absence calls and differential gene expression is presented. Expression analysis was performed using TPM and differential expression analysis was performed using the posterior probability that the gene is differentially expressed (PPDE) and posterior fold change (postFC). Genes with an average of TPM > 2 across biological replicates were compared between zones and genotypes (Figure 1; data summarized in Data File 1). Following EBSeq analysis, genes were filtered for PPDE = 1 to identify those which were differentially expressed. Of the differentially expressed genes, those with postFC (*dgt* over WT) > 2 were considered upregulated in *dgt* and those with postFC < 0.5 were considered downregulated in *dgt*. Upregulated and downregulated genes were compared between zones and genotypes (Figure 2; data summarized in Data File 2). Principal component analysis (PCA) of the data was performed using TPM values of each gene and was calculated using the scikit-learn^17^ PCA function with default parameters (Figure 3).

**Figure 1.**
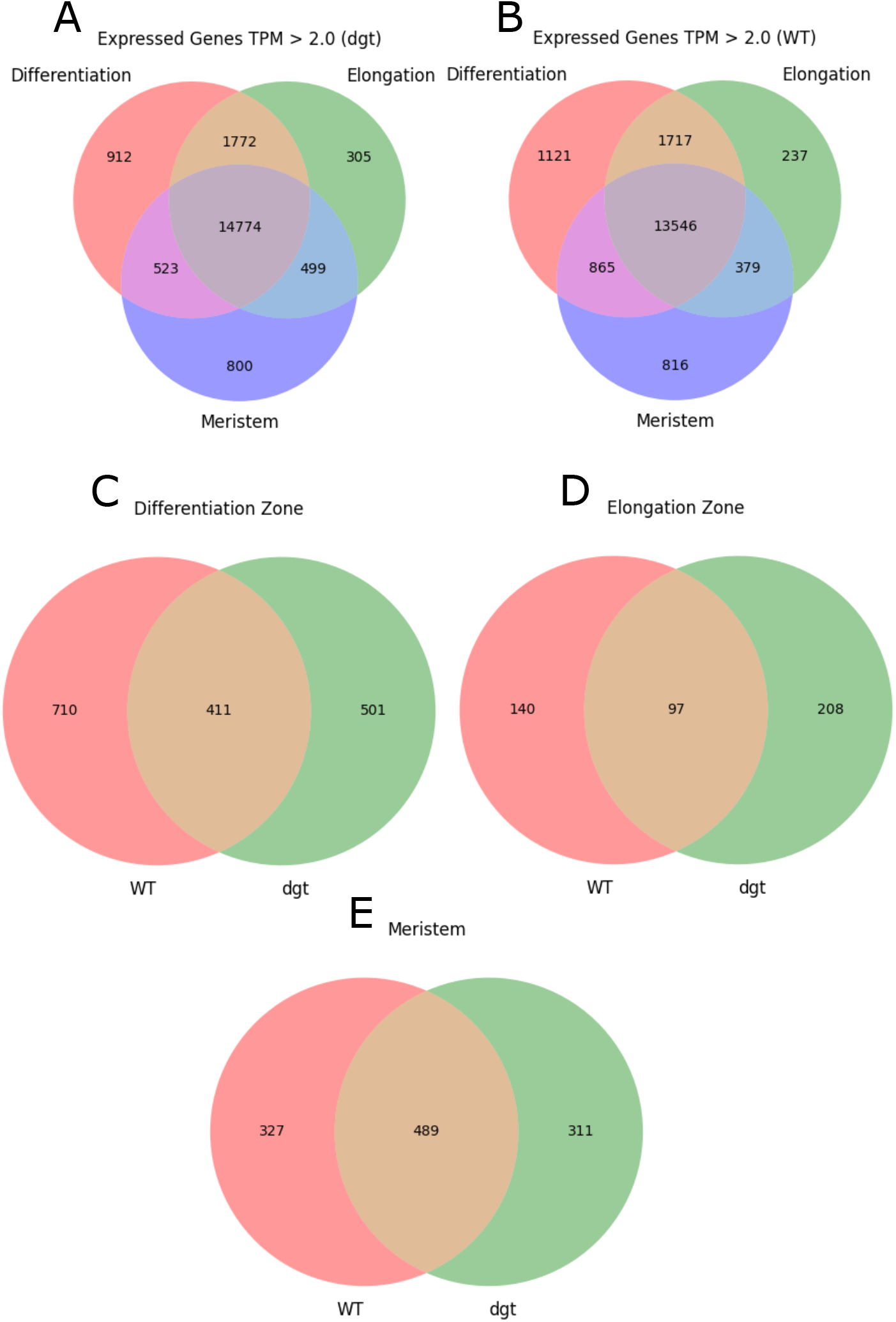
Presence/Absence Calls. (A) Number of genes expressed in each zone of *dgt* roots, TPM > 2.0. (B) Number of genes expressed in each zone of WT roots, TPM > 2.0. (C) Number of genes expressed exclusively in the differentiation zone with TPM > 2.0, compared between WT and *dgt*. (D) Number of genes expressed exclusively in the elongation zone with TPM > 2.0, compared between WT and *dgt*. (E) Number of genes expressed exclusively in the meristem with TPM > 2.0, compared between WT and *dgt*.

**Figure 2.**
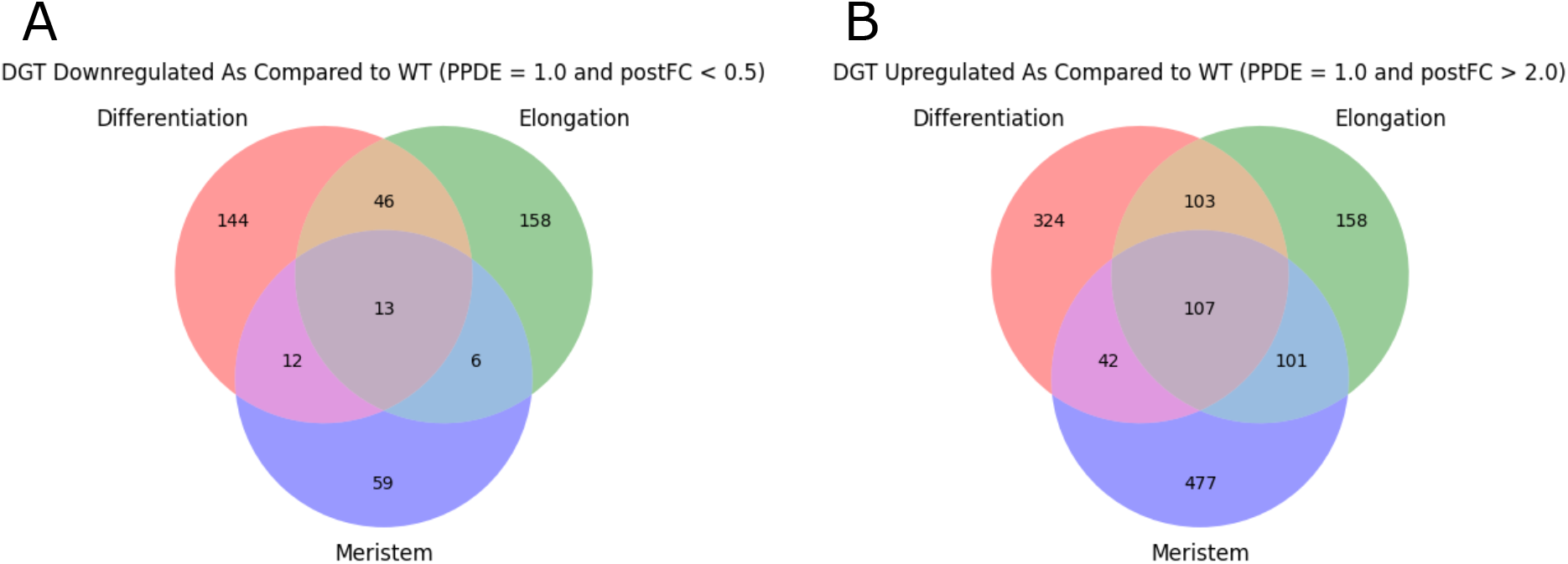
Differentially Expressed Genes. (A) Differentially expressed genes downregulated in each zone with postFC < 0.5, PPDE = 1 in *dgt* vs WT. (B) Differentially expressed genes upregulated in each zone with postFC > 2, PPDE = 1 in *dgt* vs. WT.

**Figure 3.**
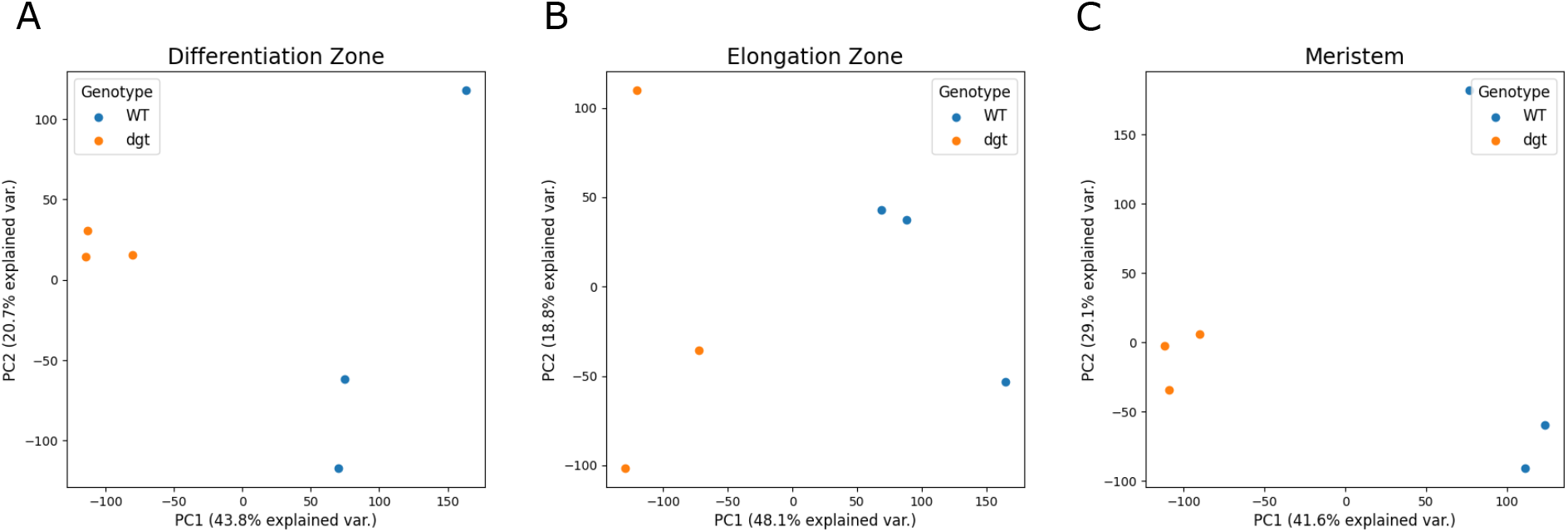
Principle Component Analysis (PCA) (A) PCA of sequenced samples in differentiation zone (B) PCA of sequenced samples in elongation zone (C) PCA of sequenced samples in meristem.

## Conclusion

It is clear from the outcomes in Figures 1 and 2 that reduced function of DGT has sweeping effects on the transcriptome in all three developmental zones examined in this study, supporting the concept that DGT very likely plays important and potentially complex roles in multiple developmental pathways. Additionally, PCA using TPM values for each sample demonstrated that while there was some variance within the replicate pools, replicates from different genotypes were distinctly separate from each other (Figure 3). Further functional genomics studies are needed to narrow down the most likely direct interactions with DGT, leading to the identification of specific functional roles. We hope that this dataset will be of value to the community in future studies in this area.

## Data Availability

Heinz 1706 genome available from ftp://ftp.ncbi.nlm.nih.gov/genomes/all/GCF/000/188/115/GCF_000188115.4_SL3.0

The raw RNA-Seq reads and expression estimates provided by RSEM were deposited into the NCBI SRA and GEO archives and will be made available upon publication, along with Data Files 1 and 2.

## Software Availability

The pipeline used to process the data will be made available upon publication.

RSEM version 1.3.1 available from https://github.com/deweylab/RSEM/releases/tag/v1.3.1 EBSeq version 1.26.0 available from https://bioconductor.org/packages/3.11/bioc/html/EBSeq.html

## Acknowledgements

We thank Jordan Holdaway for her work on the initial raw data processing scripts.

